# Accurate Phasing of Pedigree Genotypes Using Whole Genome Sequence Data

**DOI:** 10.1101/148510

**Authors:** A.N. Blackburn, M.Z. Kos, N.B. Blackburn, J.M. Peralta, P. Stevens, D.M. Lehman, L. Blondell, J. Blangero, H.H.H. Göring

**Author notes:** Corresponding author: Address: August Blackburn Ph.D., 3463 Magic Drive Suite 320, San Antonio, TX 78229.

## Abstract

Phasing, the process of predicting haplotypes from genotype data, is an important undertaking in genetics and an ongoing area of research. Phasing methods, and associated software, designed specifically for pedigrees are urgently needed. Here we present a new method for phasing genotypes from whole genome sequencing data in pedigrees: PULSAR (<u>P</u>hasing <u>U</u>sing <u>L</u>ineage <u>S</u>pecific <u>A</u>lleles / <u>R</u>are variants). The method is built upon the idea that alleles that are specific to a single founding chromosome within a pedigree, which we refer to as lineage-specific alleles, are highly informative for identifying haplotypes that are identical-by-decent between individuals within a pedigree. Through extensive simulation we assess the performance of PULSAR in a variety of pedigree sizes and structures, and we explore the effects of genotyping errors and presence of non-sequenced individuals on its performance. If the genotyping error rate is sufficiently low PULSAR can phase > 99.9% of heterozygous genotypes with a switch error rate below 1 x 10^-4^ in pedigrees where all individuals are sequenced. We demonstrate that the method is highly accurate and consistently outperforms the long-range phasing approach used for comparison in our benchmarking. The method also holds promise for fixing genotype errors or imputing missing genotypes. The software implementation of this method is freely available.

## Introduction

Haplotypes, which are combinations of alleles at different polymorphic sites occurring on the same DNA molecule, are important in the study of genetics.(Tewhey et al. 2011) Haplotypes are useful for imputation of alleles at ungenotyped loci, identification of genomic regions shared identical-by-descent, genotype error identification and correction, identification of compound heterozygosity, and analysis of parent-of-origin effects, among many other topics. Haplotypes can also be used in place of alleles at individual variable sites in association testing. Presently the most popular sequencing platforms and associated software packages report genotypes for individual polymorphisms, and thus haplotypes must be inferred algorithmically. Phasing, the process of analyzing genotypes to predict haplotypes, is therefore an important undertaking in genetics and an ongoing area of research.(Browning and Browning 2011)

While family studies have not been popular for association-based gene mapping on complex traits recently, this situation is changing with the emerging interest in rare variants, which are arguably more easily and more powerfully studied in family data. In any case, the increase in the numbers of study subjects in a population being sequenced will necessarily lead to inclusion of more - and more closely - related individuals, by chance. For phasing, pedigrees provide additional information compared to unrelated individuals (which are in reality distantly related). The direct observation of inheritance of alleles from one generation to the next, which is possible in families, can be used to establish phase empirically, and hence families conceptually allow for highly accurate estimation of haplotypes. This also means that phasing of family data does not necessarily have to bear the computational expense of dealing with a vast number of additional samples in the form of reference panels, and that family data can be useful in populations for which good reference panels are not available. However, pedigree data brings with it substantial and unique computational challenges. Thus, computational methods and ready-to-use software tailor-made for phasing related individuals in pedigrees using whole genome sequence data are needed.

Here we describe a novel fast algorithm, which we have named PULSAR (<u>P</u>hasing <u>U</u>sing <u>L</u>ineage <u>S</u>pecific <u>A</u>lleles / <u>R</u>are variants), that phases whole genome sequencing data in families. PULSAR is built upon the idea that alleles that are specific to a single founding chromosome within a pedigree, which we refer to as lineage-specific alleles (LSAs), are highly informative for identifying haplotypes that are identical-by-decent (IBD) between individuals within the pedigree. In humans, each chromosome carries many such rare variants, which we can now interrogate with whole genome sequencing, allowing for reasonably dense haplotype maps comprised of LSAs, from which we can observe inheritance of haplotypes empirically. This initial haplotype map can then be expanded to include non-LSAs, which tend to be more common variants. We demonstrate the utility of PULSAR and its software implementation on simulated data as well as real whole genome sequencing data and compare its performance to long-range phasing.

## Results

### Description of method

The general steps of the PULSAR algorithm are as follows: 1) identify alleles that are likely to be lineage-specific (i.e., LSAs); 2) identify haplotypes, their boundaries, and their inheritance using LSAs; 3) extend the estimated boundaries of these haplotypes based on the observation that individuals will share at least one allele at loci where they share a haplotype IBD (i.e., the idea behind long range phasing); and 4) assign alleles to the haplotypes. Our method assumes that the physical location of variants is known, and optionally makes use of external information regarding allele frequencies. Below, we will describe these steps in more detail.

#### Identifying putative lineage-specific alleles

The premise of the PULSAR algorithm is that when an allele is present in a single chromosome among the founders of a given pedigree, then any direct descendent of that founder also carrying the allele can be assumed to share the chromosomal segment harboring that locus IBD, which allows for empirical estimation of the inheritance of haplotypes within the pedigree. Exceptions to this logic, such as *de novo* mutation, are real but infrequent enough not to undermine the premise of the approach. Since not all founders will be sequenced in many pedigrees, an initial challenge is identifying those alleles that are specific to a single founding chromosome. We identify potential LSAs by analyzing the pattern of individuals within a pedigree carrying a given allele. The implementation of this approach is simple; we search for alleles for which all individuals carrying the allele share at least one founder within the known structure of the pedigree under consideration. We check that the direct lineage between each founder and each person carrying the allele also carries the allele (at least for those individuals that are sequenced) and that the allele is not homozygous in any individuals. For now, the algorithm does not accommodate the presence of inbreeding loops, wherein LSAs could be homozygous in inbred individuals. Figure 1A shows an example of a family where individuals carry an allele that cannot be lineage-specific since the carriers do not share a common founder. Figure 1B shows an example of a family where individuals carry a putative lineage specific allele. Note that at this point we are only seeking to identify *putative* LSAs.

**Figure 1.**
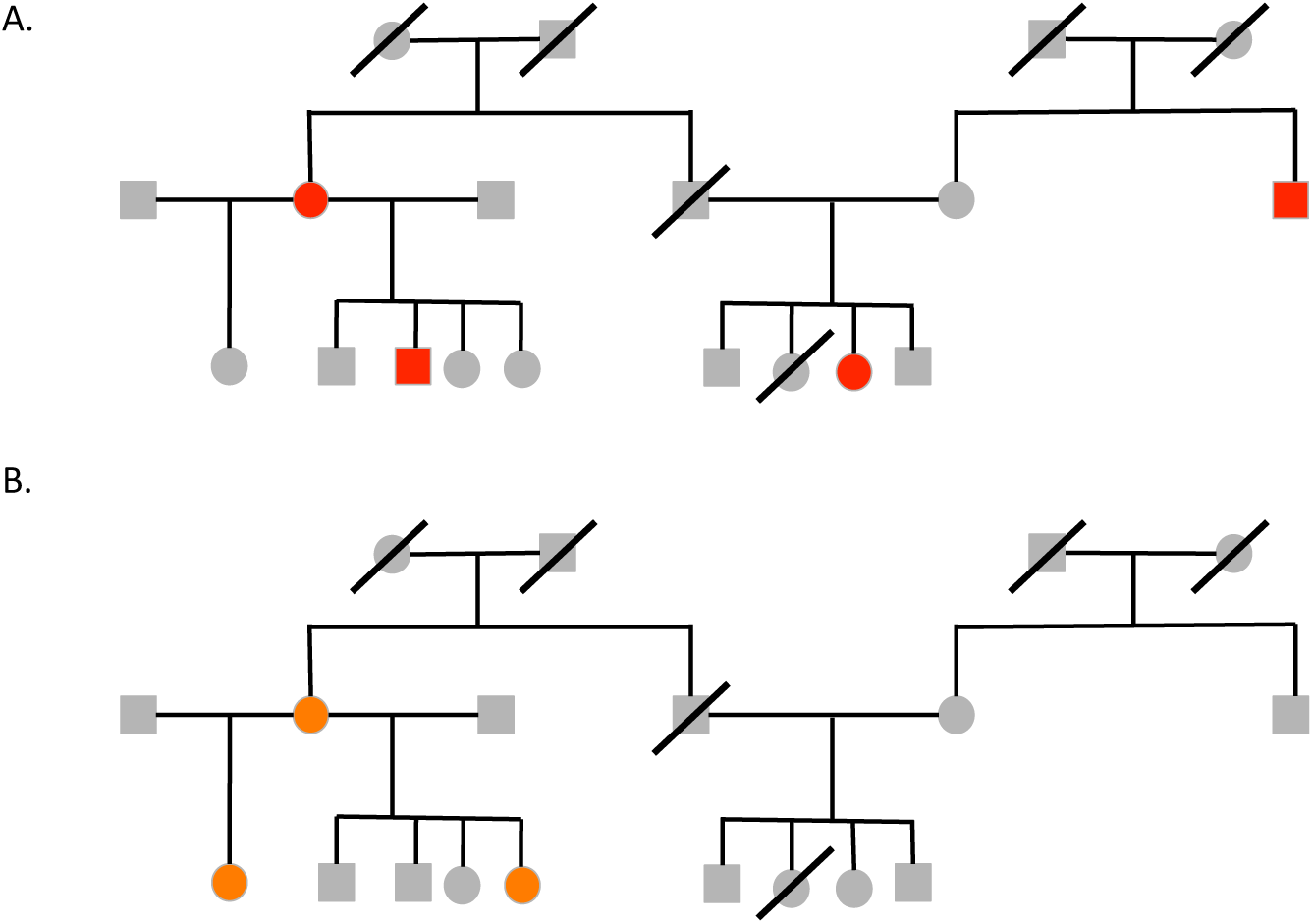
Inheritance patterns of lineage specific alleles. Panel A shows an example of a pattern of individuals carrying an allele in which the allele cannot be lineage-specific since individuals carrying the allele do not share a common founder. Panel B shows an example where the allele could be lineage specific. The individuals carrying the allele have two common founders.

#### Constructing haplotypes boundaries and establish haplotype inheritance using LSAs

Using the set of putative LSAs from the previous step, we then seek to establish boundaries within which haplotypes are shared IBD between a set of related individuals in a pedigree. Note that at this point the putative LSAs may include some false positives. However, for now, let us assume that we are dealing only with true LSAs, and relax this assumption later. The set of individuals carrying a true LSA will share an allele IBD at nearby loci, assuming absence of meiotic recombination between the loci, mutation, or genotyping error. Conversely, a change in pattern (of which individuals share a LSA) from one locus to the next indicates a recombination event(s) within the meioses that give rise to the haplotypic lineage. Hence, by tracking which individuals share neighboring LSAs along the chromosome we can establishes the boundaries of a given haplotype. While the concept is straightforward, complications arise because at any given region of the diploid genome all individuals carry two haplotypes, one maternal and one paternal. Therefore, it is necessary to track two separate haplotypes simultaneously. We implement this approach using a rules-based algorithm to identify the changes in the patterns of LSA sharing along a chromosome.

If not all individuals are sequenced in a given pedigree, which is often the case, there will likely be some false positives among the putative LSAs identified in the previous step. In other words, some of the putative LSAs are in reality not IBD but merely identical-by-state (IBS). To reduce the risk of mistaking IBS for IBD, and thus inferring wrong haplotypes, we require a predetermined number of neighboring putative LSAs to be shared in order to demarcate a new haplotype. A very low number of changed LSA pattern observations are required to infer, with high confidence, that there is a true recombination event because it is highly unlikely that the same individuals will share putative LSAs over a given region of a chromosome if the genotypes are a product of chance (such as being IBS or perhaps due to genotyping error) and not truly lineage specific. Here we set this heuristic to 5 observations of neighboring putative LSAs with the same changed pattern of individuals sharing the putative LSAs.

After identifying the haplotypes a proband carries along the chromosome, we perform two procedures to establish chromosomal length haplotypes from these smaller haplotype segments. First, during the course of identifying the haplotypes if a change is observed in one set of individuals (one haplotype segment changes) we assume that the previous and new haplotype segments are on the same chromosome in the person where the pattern has changed. Second, for non-founders with a sufficient number of informative relatives with sequencing data we identify if the haplotype is shared with the maternal or paternal side of the proband’s relatives. Since each proband inherits one chromosome from each parent, haplotypes that are shared with either paternal or maternal relatives are assumed to be on either the paternally or maternally inherited chromosome, respectively.

#### Extension of haplotype boundaries using IBS allele sharing

While the human genome contains a vast number of rare variants, many of which will be lineage-specific in a given pedigree, the density of LSAs nonetheless limits the precision with which the boundaries of haplotypes can be determined. We extend the haplotype boundaries observed in a proband with a simple heuristic, namely that individuals sharing a haplotype must share an allele IBD (and thus also IBS) at each locus in the haplotype. This same heuristic is central to the rationale behind long-range phasing, and our implementation is similar in that we search for opposing homozygous genotypes. The primary difference is that we are not using this rationale to discover shared haplotypes, rather only extending predetermined haplotypes, carried by known individuals, for relatively small unresolved segments of the genome (typically <1% of the genome; we present an investigation of the density of LSA coverage below). This haplotype extension step proceeds in both directions into the unassigned gap between two neighboring haplotypes identified in the previous step, and we extend haplotypes only so far that there is no overlap.

#### Mapping alleles to haplotypes

After establishing the haplotype boundaries and inheritance throughout the pedigree, we then map alleles for all variants onto these haplotypes. This step could be integrated into the prior steps, but in our implementation we assign alleles to haplotypes (including LSAs) as a separate, final step of the procedure. Homozygous genotypes are straightforward to map onto haplotypes, with each of the two haplotypes carrying the same allele; this is done first. Considering each variant independently, we then phase heterozygous genotypes in individuals for which at least one allele has already been mapped onto one of the two haplotypes they carry at that genomic location. We repeat this mapping process iteratively, now incorporating the mapped alleles from heterozygous genotypes from the previous iteration, until no additional genotypes can be phased. When all individuals within a pedigree are sequenced, and all of the haplotypes each individual carries are known, barring genotyping errors this procedure can resolve the phase of all combinations of genotypes with exception of the case where every individual is heterozygous, a scenario that becomes less likely in larger pedigrees.

PULSAR considers evidence from multiple individuals (if multiple people carry the haplotype) when assigning alleles to haplotypes. In the presence of genotyping errors it becomes possible for the genotypes of multiple individuals to provide conflicting support for which allele is carried on a given haplotype. As an example, two individuals share a haplotype but have opposing homozygous genotypes. In these ambiguous cases, when more than one allele is supported by the inferred haplotype sharing and individual genotypes, the allele supported by the majority of individuals is assigned to that haplotype. When the correct alleles are assigned to both haplotypes carried by an individual, reconstructed genotypes from these haplotypes can be used to correct genotyping errors. However, in some cases there is no majority of individuals with which PULSAR can correct genotyping errors. As an example, in trios no haplotype is shared between more than two people, and thus no majority can be formed in order to correct genotyping errors. This same principle is true for haplotypes that are shared between only two or fewer individuals within larger pedigrees, such as haplotypes shared between married-in founders and only one offspring when the lineage is not passed on to subsequent generations.

## Benchmarking

### Coverage and allele-frequency distribution of lineage-specific alleles

As we described, the PULSAR algorithm is based on the idea that variants that are introduced into a pedigree via a single founding chromosome, i.e. LSAs, can be used as convenient tags to trace the inheritance of haplotypes within pedigrees. For this approach to be practical, it is crucial that LSAs exist in sufficient density in realistic pedigree structures. Thus, we have estimated the degree of LSA coverage using real sequencing data with known phase, namely the X chromosomes from the British (GBR) and Finnish (FIN) cohorts from the 1000 Genomes Project(Genomes Project et al. 2015). If we view pedigree founders as a random population sample, then various key aspects of LSA coverage can easily be determined by permutation for different pedigree structures. Supplemental Figure 1 shows the distribution of the number of LSAs per Mb along the X chromosome as a function of the number of pedigree founders for the GBR and FIN cohorts. In British, the density ranges from a median of 135.4 LSAs per Mb in 2-founder pedigrees (median inter-LSA distance of 874 bp, with a maximum of 5.14Mb which includes a gap in coverage caused by the centromere) to 14.8 LSAs per Mb with 15 founders (median distance of 19.0Kb, with a maximum of 5.39Mb which includes a gap in coverage caused by the centromere). We estimate that a pedigree with 170 founders would have ~1.0 LSAs per Mb. In Finns, a population with a lower genetic diversity (primarily due to a small founding population), there are comparatively fewer LSAs, a median of 133.1 LSAs per Mb in 2-founder pedigrees and 13.4 LSAs per Mb with 15 founders. The trend to fewer LSAs in larger pedigrees make sense since our definition of LSA is based on a single founding occurrence per pedigree, and thus fewer polymorphic sites qualify the more founders a pedigree contains.

With regard to the allele frequency distribution of LSAs, since the probability of a single founding event in a pedigree depends on the population prevalence of an allele, rare alleles are more likely to be specific to a single founding chromosome, and the enrichment for rare alleles among all LSAs will be greater in pedigrees with more founders. This is what we observe. In British, with 2 founders ~13% of LSAs have minor allele frequencies <5% (median allele frequency is 21.7%), whereas with 15 founders ~91% of LSAs have minor allele frequencies <5% (median allele frequency is 2.2%). These observations indicate that MAF estimates, either from the dataset in hand or from a reference panel, are expected to be a useful filter for decreasing the false positive rate when identifying LSAs is ambiguous (perhaps due to non-sequenced individuals). In any case, the important take-home message with regard to our phasing method is that in humans there appear to be sufficiently many LSAs to make them useful as a starting point for phasing of WGS data in pedigrees.

### Pedigrees with complete sequencing

We measured the performance of PULSAR and AlphaPhase1.1 in pedigrees varying in size and structure under the ideal scenario in which all individuals in the dataset are sequenced without genotyping errors. Table 1 presents a comparison of the SER and the percentage of heterozygous markers phased for these simulations. Across all simulations, PULSAR produced lower SERs and a higher percentage of heterozygous markers phased than AlphaPhase1.1. In the case of nuclear pedigrees some markers were heterozygous in all individuals in a pedigree, a situation that is unresolvable without external data. Excluding these unresolvable cases, the observed number of phased heterozygous genotypes was >99.9% of the theoretical upper limit using PULSAR.

**Table 1.** Complete and Accurate Data

| Pedigree Size | Founders | Generations | Switch Error Rate |  | Percentage of Heterozygous Genotypes Phased |  |
| --- | --- | --- | --- | --- | --- | --- |
|  |  |  | PULSAR | AlphaPhase1.1 | PULSAR | AlphaPhase1.1 |
| 3 | 2 | 2 | 0 | $4.17 \times 10^{-2}$ | 0.8164 | 0.7425 |
| 4 | 2 | 2 | 0 | $8.46 \times 10^{-2}$ | 0.8499 | 0.8193 |
| 5 | 2 | 2 | $1.04 \times 10^{-5}$ | $9.20 \times 10^{-2}$ | 0.9420 | 0.8994 |
| 6 | 2 | 2 | $1.28 \times 10^{-5}$ | $8.44 \times 10^{-2}$ | 0.9887 | 0.9312 |
| 7 | 2 | 2 | $1.79 \times 10^{-5}$ | $8.93 \times 10^{-2}$ | 0.9998 | 0.9398 |
| 8 | 2 | 2 | $1.56 \times 10^{-5}$ | $9.28 \times 10^{-2}$ | 0.9999 | 0.9514 |
| 9 | 2 | 2 | $1.68 \times 10^{-5}$ | $7.84 \times 10^{-2}$ | 0.9999 | 0.9464 |
| 11 | 5 | 3 | $1.90 \times 10^{-5}$ | $5.52 \times 10^{-2}$ | 0.9996 | 0.9794 |
| 14 | 6 | 4 | $4.85 \times 10^{-5}$ | $5.46 \times 10^{-2}$ | 0.9979 | 0.9815 |
| 20 | 6 | 4 | $1.07 \times 10^{-4}$ | $4.10 \times 10^{-2}$ | 0.9966 | 0.9846 |
| 28 | 8 | 4 | $3.43 \times 10^{-5}$ | $4.73 \times 10^{-2}$ | 0.9989 | 0.9849 |
| 55 | 17 | 6 | $1.15 \times 10^{-4}$ | $5.50 \times 10^{-2}$ | 0.9989 | 0.9747 |
| 78 | 21 | 6 | $7.15 \times 10^{-4}$ | $5.53 \times 10^{-2}$ | 0.9973 | 0.9744 |
| 94 | 28 | 5 | $2.64 \times 10^{-4}$ | $6.26 \times 10^{-2}$ | 0.9966 | 0.9696 |

### Effect of genotyping errors

We sought to assess the robustness of PULSAR to genotyping errors. Table 2 presents a comparison of the switch error rate and the percentage of heterozygous markers phased for simulations wherein the genotyping accuracy is 99.0% (assuming, for simplicity, an equal error rate for all variants). Again, PULSAR produced lower switch error rates and higher percentage of heterozygous markers phased than AlphaPhase1.1.

**Table 2.** Genotyping Errors

| Pedigree Size | Founders | Generations | Switch Error Rate |  | Percentage of Heterozygous Genotypes Phased |  |
| --- | --- | --- | --- | --- | --- | --- |
|  |  |  | PULSAR | AlphaPhase1.1 | PULSAR | AlphaPhase1.1 |
| 9 | 2 | 2 | $1.74 \times 10^{-3}$ | $7.22 \times 10^{-2}$ | 0.9997 | 0.8960 |
| 11 | 5 | 3 | $8.33 \times 10^{-3}$ | $6.95 \times 10^{-2}$ | 0.9961 | 0.8800 |
| 14 | 6 | 4 | $8.44 \times 10^{-3}$ | $6.22 \times 10^{-2}$ | 0.9837 | 0.9093 |
| 20 | 6 | 4 | $6.99 \times 10^{-3}$ | $5.33 \times 10^{-2}$ | 0.9830 | 0.9316 |
| 28 | 8 | 4 | $5.95 \times 10^{-3}$ | $6.36 \times 10^{-2}$ | 0.9912 | 0.9242 |
| 55 | 17 | 6 | $7.51 \times 10^{-3}$ | $6.71 \times 10^{-2}$ | 0.9629 | 0.9149 |
| 78 | 21 | 6 | $9.28 \times 10^{-3}$ | $6.73 \times 10^{-2}$ | 0.9437 | 0.9228 |
| 94 | 28 | 5 | $8.41 \times 10^{-3}$ | $7.60 \times 10^{-2}$ | 0.9026 | 0.9091 |

For the case of nuclear pedigrees, we created multiple simulated genotyping datasets with a range of genotyping accuracies (from 99.0% to 100.0%). Genotyping errors were simulated for 20 datasets at each genotyping accuracy level for each pedigree. As shown in Figure 2, as the genotyping error rate increases, the switch error rate increases linearly in the simulated pedigrees. Genotyping errors had an attenuated negative affect on switch error rate in the pedigrees with more children. The reason being that in a larger pedigree, more individuals potentially share genomic sections IBD with one another, enabling PULSAR to identify haplotypes more reliably in the presence of genotyping errors.

**Figure 2.**
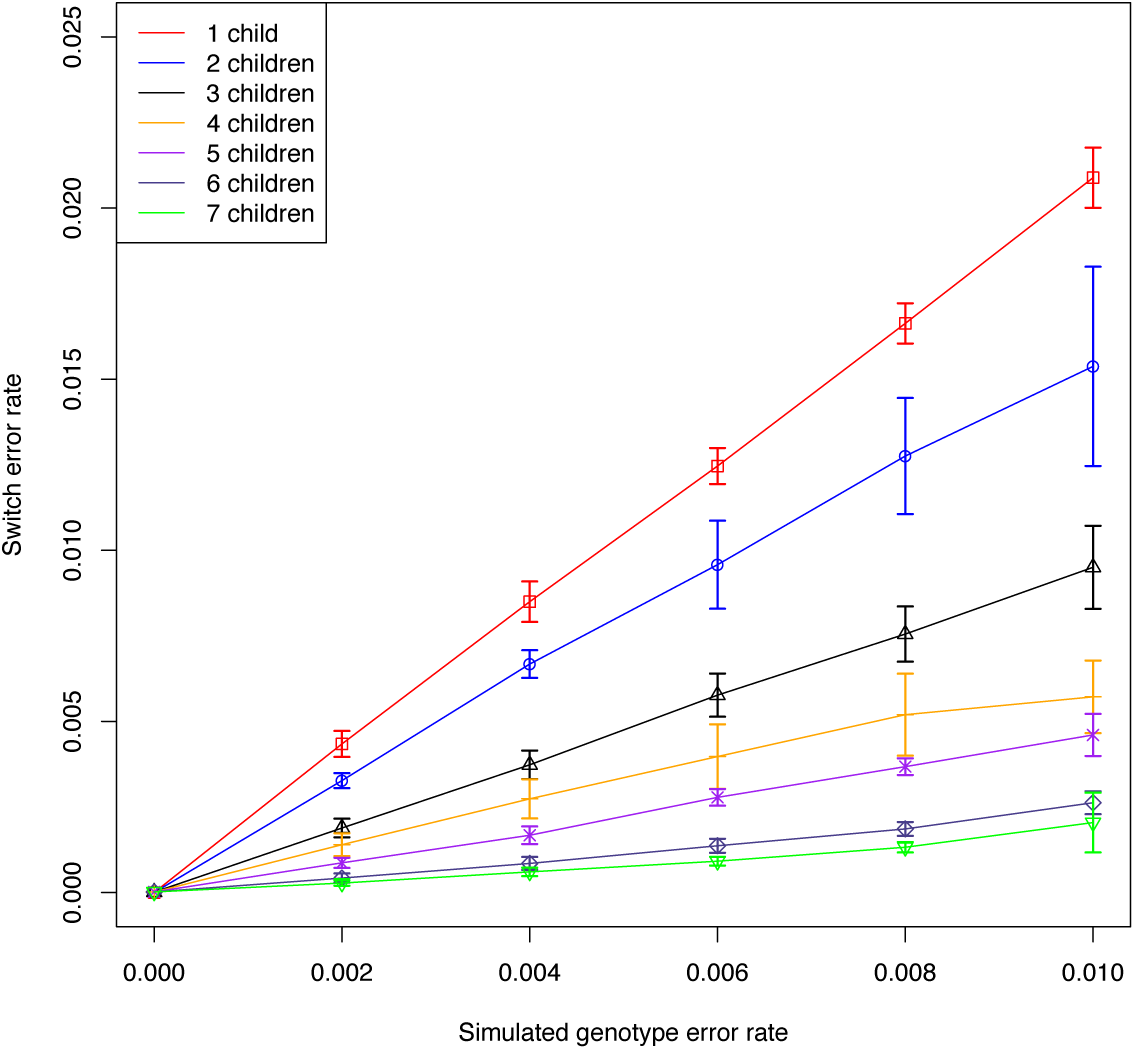
Effect of genotyping accuracy and IBD sharing on switch error rate. The plot shows the switch error rates (with estimated error bar) from 20 simulations at varying genotyping accuracies in nuclear pedigrees with a range of 1-7 children, in which each pedigree with fewer children is a subset of the pedigrees with more children. Genotyping errors increase the switch error rate in a predictable manner. Increased IBD sharing within the pedigree attenuates this effect.

Based on the haplotypes output by PULSAR we reconstructed genotypes and compared them to the true genotypes (i.e., without introduced genotyping errors). As shown in Figure 3, the accuracy of the reconstructed genotypes decreases linearly with the simulated error rate across pedigrees. The reconstructed genotypes accuracies from the trio pedigree are very similar to the accuracy of the simulated genotypes, which is expected because the pedigree structure lacks a majority of haplotypes with which to correct genotyping errors (as outlined previously). In fact, the reconstructed genotype error rate is slightly higher than the true genotype error rate, because the algorithm introduces some (though very few) errors when haplotype boundary estimation is imperfect. However, PULSAR is able to correct genotypes in pedigrees with 2 or more children, performing better in pedigrees with more children. Together these observations also demonstrate that increased IBD sharing between pedigree members improves the performance of the PULSAR algorithm.

**Figure 3.**
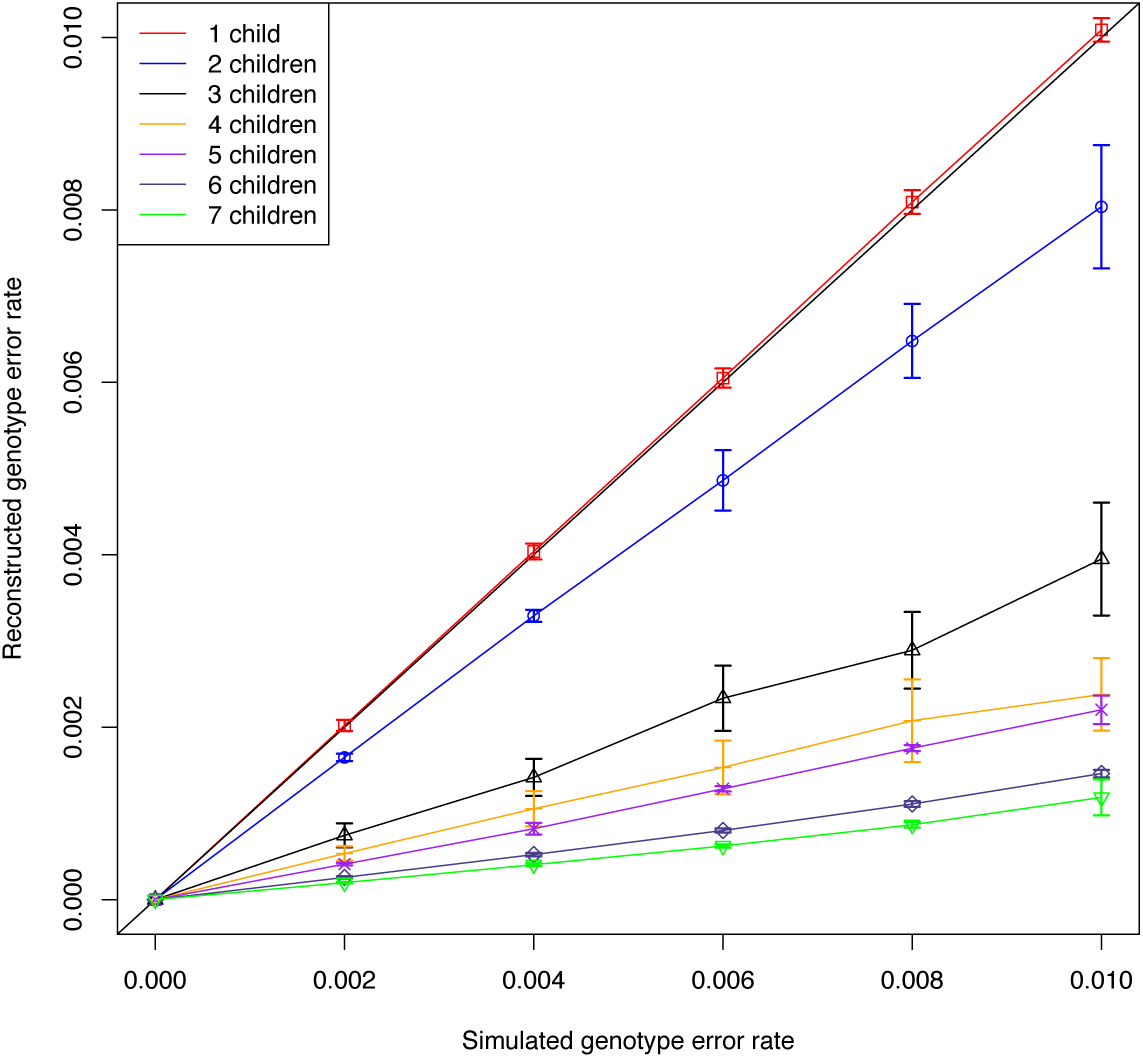
Effect of IBD sharing on correcting genotypes through imputation. The plot shows the accuracies (and associated estimated error) of genotypes reconstructed from haplotypes produced by PULSAR for 20 simulations at varying genotyping accuracies in nuclear pedigrees with a range of 1-7 children, in which each pedigree with fewer children is a subset of the pedigrees with more children. The diagonal is present to show the equivalence between the axes. Increased IBD sharing improves the ability of the PULSAR algorithm to correct genotypes through imputation.

### Effect of ungenotyped individuals

Ungenotyped individuals in a pedigree are another frequent complication in real datasets. PULSAR will not phase heterozygous genotypes for individuals in genomic regions where the individual does not share a chromosomal segment IBD with another sequenced individual, unless done erroneously after falsely inferring IBD sharing. Non-sequenced individuals can affect the false positive rate among putative LSAs identified by PULSAR, which will ultimately decrease the accuracy of haplotype boundary and sharing estimation.

Table 3 presents a comparison of the switch error rate and the percentage of heterozygous markers phased for simulations wherein some individuals are missing sequencing data. For the SAMAFS pedigrees, individuals with real sequencing data were given sequencing data in the simulation. We performed two simulations involving the nuclear family, wherein one and both parents were not sequenced. The comparative estimates for the scenario of complete genotyping of every pedigree member are already shown in Table 1. The number of ungenotyped individuals and, more importantly, the location of the ungenotyped individuals within the pedigree have notable effects on the haplotype phasing results. In the case of the nuclear families, children missing sequencing data have only a small negative effect on the accuracy of PULSAR, as this situation does not lead to an increase in the number of falsely inferred LSAs, though the number of observations of each haplotype is reduced on average. Children in a nuclear family missing sequencing data are essentially the same as reducing the size of the nuclear family by the number of unsequenced children. A consequence is that a higher portion of variants will be heterozygous in all sequenced individuals (this is only of practical importance if the number of sequenced individuals in a pedigree is small). For that reason, the main impact of missing children in nuclear families is a lowered percentage of loci that are phased because PULSAR does not resolve cases wherein all individuals are heterozygous (see the estimates from Table 1 for nuclear families of different sizes). The effect of missing parents is much greater because this scenario leads to ambiguity in identifying LSAs. In the simulated pedigree with one missing parent the false positive rate among putative LSAs was 15.9% compared to 0% when both parents are sequenced, and the SER increased from 1.7x10^-5^ to 4.0x10^-2^. The total number of heterozygous markers phased dropped from 99.99% to 99.45%.

**Table 3.**
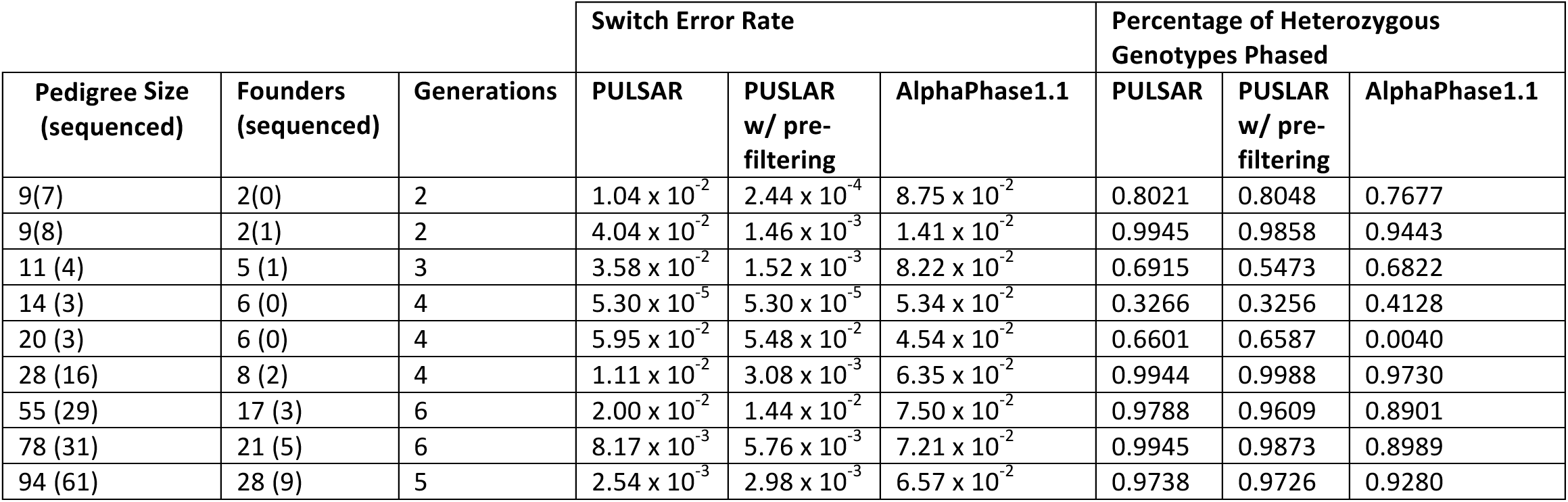
Missing Individuals

### Mitigating the negative effect of ungenotyped individuals

We sought to mitigate the negative effect of ungenotyped individuals on the performance of PULSAR. We investigated the extent to which filtering LSAs based on minor allele frequency would help. Table 3 also presents a comparison of the effects of missing individuals in nuclear families wherein we have pre-filtered putative LSAs based on a minor allele frequency of <5%. In the nuclear pedigree with 7 children with one non-sequenced parent, the rate of falsely inferred LSAs was 15.9% prior to filtering and 1.50% afterward. The SER dropped from 4.0 x 10^-2^ to 1.5 x 10^-3^. Pre-filtering is not without cost, however. The total number of putative LSAs dropped from 61,160 to 7,850. The total percentage of heterozygous markers phased dropped from 99.45% to 98.58%. In the nuclear family, considering the tradeoffs between the accuracy of the phasing results (measured using SER) and the percentage of heterozygous markers phased, filtering based on MAF performed better, improving the overall number of heterozygous markers phased correctly, which can be measured by the product of the accuracy (1-SER) and the percentage of heterozygous markers phased.

In the largest pedigree in this study, wherein 61 of 94 (64.9%) individuals have sequencing data, filtering using MAF <5% had only a minor effect on the results. The false positives among putative LSAs dropped from 0.0099 to 0.0091, the total number of putative LSAs dropped from 44889 to 40060, the SER went up from 2.54x10^-3^ to 2.98 x10^-3^, and the percentage of heterozygous variants phased dropped from 97.4% to 97.3%. In pedigrees with many founders, the putative LSAs that PULSAR identifies already have low allele frequencies for the most part, and hence filtering at a 5% prevalence threshold has little effect. It is possible that an even lower MAF filter would be advantageous, though it is clear that MAF filtering is more advantageous in smaller pedigrees with fewer founders.

Non-sequenced individuals have a negative effect on the performance of the PULSAR algorithm, but not in all cases. Since it is not always straightforward to estimate which cases of missing individuals are detrimental to the effectiveness of the PULSAR algorithm we have created a tool that uses a Monte-Carlo gene-dropping approach to estimate the expected percentage of the genome that each individual will share at least one haplotype IBD with another sequenced individual in the pedigree. This tool can be used for *a priori* estimation of the applicability of the PULSAR algorithm to a pedigree dataset given the sequenced individuals within the pedigree.

### Performance on data with sequencing errors and missing individuals

We then sought to benchmark our approach in realistic data, having both missing individuals and genotyping errors. To do this we only examined genotypes for individuals with whole genome sequencing data available in the SAMAFS. The nuclear pedigree was set up so that genotypes for one parent were missing. We simulated genotypes with a conservative genotype accuracy of 99.0%. Most sequencing platforms outperform this benchmark easily with appropriate read depth, and thus 99.0% accuracy is conservative. Results for these simulations are presented in Table 4. With few exceptions, PULSAR outperformed AlphaPhase1.1 both in accuracy and in completeness.

**Table 4.** Realistic Data

| Pedigree Size<br>(sequenced) | Founders<br>(sequenced) | Generations | Switch Error Rate |  | Percentage of Heterozygous<br>Genotypes Phased |  |
| --- | --- | --- | --- | --- | --- | --- |
|  |  |  | PULSAR w/<br>pre-filtering | AlphaPhase1.1 | PULSAR w/<br>pre-filtering | AlphaPhase1.1 |
| 9 (8) | 2 (1) | 2 | $1.85 \times 10^{-2}$ | $1.32 \times 10^{-1}$ | 0.9370 | 0.8879 |
| 11 (4) | 5 (1) | 3 | $1.85 \times 10^{-2}$ | $6.24 \times 10^{-2}$ | 0.5207 | 0.5331 |
| 14 (3) | 6 (0) | 4 | $1.96 \times 10^{-2}$ | $5.71 \times 10^{-2}$ | 0.3327 | 0.3166 |
| 20 (3) | 6 (0) | 4 | $7.25 \times 10^{-2}$ | * | 0.6569 | * |
| 28 (16) | 8 (2) | 4 | $7.71 \times 10^{-2}$ | $7.50 \times 10^{-2}$ | 0.9617 | 0.9026 |
| 55 (29) | 17 (3) | 6 | $3.83 \times 10^{-2}$ | $8.38 \times 10^{-2}$ | 0.8734 | 0.8155 |
| 78 (31) | 21 (5) | 6 | $1.93 \times 10^{-2}$ | $8.22 \times 10^{-2}$ | 0.9274 | 0.8295 |
| 94 (59) | 28 (9) | 5 | $1.45 \times 10^{-2}$ | $7.83 \times 10^{-2}$ | 0.8908 | 0.8393 |

**\*Not phased successfully**

### Application to real whole genome sequencing data

Lastly, we sought to investigate the performance of PULSAR when applied to real whole genome sequencing data. To do this we used whole genome sequencing genotypes for chromosome 21 from the San Antonio Mexican American Family Study, focusing on the same multigenerational pedigrees that we have used as template pedigree structures in our simulations. This dataset contains all problems and errors inherent with real world data, including ungenotyped individuals, some missing (uncalled) genotypes within sequenced individuals, and genotyping errors. (Gross errors in pedigree relationships had been previously detected and corrected, however, but some low level of kinship between presumed unrelated founders remains possible.) The downside of real data is that the true state, here mainly phasing, is unknown. To permit a more complete characterization of the performance of the PULSAR algorithm on the real data, we created artificial blanks by masking genotypes randomly each with a probability of 1/1000, permitting us to compare the original genotype calls and the imputed genotypes after phasing. We then applied PULSAR to the manipulated WGS genotypes. We reconstructed genotypes from the phased haplotypes produced by PULSAR, and then compared them to the original genotypes. Table 5 summarizes the results for this phasing and genotype imputation experiment. PULSAR was able to phase the vast majority of observed genotypes (>97% in all pedigrees, except in one of the sparsely genotyped pedigrees). And the genotypes reconstituted from phased haplotypes matched the observed, unblanked genotypes well (>98.5% concordance). For the masked genotypes, the concordance rate between the imputed genotypes and the masked genotypes ranged from 95.5% to 98.6% across pedigrees, indicating a high degree of phasing accuracy and genotype imputation. However, only a fairly small percentage of the masked genotypes were imputed (<50%). There were also some actual missing genotypes in the original sequencing data. There is a notable difference between the percentages of masked genotypes and truly missing genotypes PULSAR was able to impute. This difference is likely due to the nature of how the missing genotypes are distributed. The masked genotypes were distributed randomly, whereas the truly missing genotypes were more likely to be found at the same marker in multiple individuals. This could be due to any number of reasons, with one reasonable hypothesis being that indels or copy number variants disrupt the diploid state of these markers in a subset of related individuals. In summary, PULSAR phased the vast majority of the observed genotypes accurately.

**Table 5.** Real Data

| Pedigree Size (sequenced) | Founders (sequenced) | Generations | Concordance rate between genotypes reconstructed from phasing results and non-masked genotypes | Percentage of non-masked genotypes phased | Concordance rate between genotypes reconstructed from phasing results and masked genotypes (ie. imputed genotypes) | Percentage of masked genotypes phased/imputed | Percentage of missing genotypes imputed |
| --- | --- | --- | --- | --- | --- | --- | --- |
| 11 (4) | 5 (1) | 3 | 0.9948 | 0.9769 | 0.9813 | 0.1853 | 0.0714 |
| 14 (3) | 6 (0) | 4 | 1 | 0.9297 | * | 0 | 0 |
| 20 (3) | 6 (0) | 4 | 0.9859 | 0.9950 | 0.9545 | 0.4221 | 0.2025 |
| 28 (16) | 8 (2) | 4 | 0.9899 | 0.9937 | 0.9798 | 0.4385 | 0.3396 |
| 55 (29) | 17 (3) | 6 | 0.9873 | 0.9885 | 0.9689 | 0.2450 | 0.2074 |
| 78 (31) | 21 (5) | 6 | 0.9943 | 0.9909 | 0.9858 | 0.3116 | 0.2603 |
| 94 (61) | 28 (9) | 5 | 0.9941 | 0.9858 | 0.9823 | 0.2866 | 0.1437 |

**\*No genotypes were imputed**

### Computation time

We sought to estimate the scalability of our software to larger pedigrees. We simulated pedigrees with two parents and a range of children between 1 and 1000. This design allows us to investigate the effect of increasing the IBD sharing within the pedigree. It appears that the computation time can be modeled well using a second order polynomial based on the number of individuals in the pedigree. We estimate, based on the estimated polynomial function, that a pedigree with 2 parents and 1000 children would take roughly 5.5 days to run on a MacBook Pro laptop with a 2.9 GHz Intel Core i7 processor. We also simulated pedigrees in which each generation consisted of one offspring of the previous generation and one married-in founder. This pedigree design was chosen because the IBD sharing within the pedigree does not increase over generations. We simulated pedigrees in the range of 2-50 generations in this manner. A linear model appears to be the best fit to describe the relationship between the number of individuals in the pedigree and the time PULSAR took to analyze the genotypes created in each simulation. We estimate that a pedigree with 500 generations and 999 individuals would take roughly 88 minutes to run.

Thus, taken together, these results show that both pedigree size and pedigree structure affect run time of the PULSAR algorithm, with a portion of the increase in pedigree size scaling linearly.

In the runs on the simulated data in the largest pedigree (94 individuals), AlphaPhase (93 seconds) was faster than PULSAR (938 seconds) on a server with 2.40 GHz Intel Xeon Core i7 CPUs running CentOS Linux 7. Although computation times become important with increasing volumes of data, we found PULSAR to be sufficiently fast for practical use on whole genome sequencing in the size of pedigrees presented here.

## Discussion

Accurate phasing information is an essential step in many genetic studies. Nonetheless, there are currently few tools designed specifically to phase genotypes in a broad range of pedigree sizes and structures, with most available methods working only in unrelated individuals or in nuclear families. In the benchmarking studies we have undertaken, PULSAR outperformed long-range phasing (using the software AlphaPhase1.1), produced low switch error rates, and phased a high percentage of heterozygous variants.

While the method clearly has merit, it has some shortcomings and limitations. In our simulations we have explored the impact of non-sequenced individuals and genotype errors, both of which are unavoidable in real datasets and weaken the performance of PULSAR, as well as alternative approaches for phasing. From our simulations of realistic data, we examined some pedigrees with very sparse sequencing, such as 3 out of 20 individuals being sequenced. Even in these problematic pedigrees, with many missing individuals, PULSAR performed fairly well, erring on the side of phasing fewer heterozygous markers rather than phasing them incorrectly. The percentage of alleles individuals share IBD with at least one other sequenced individual is the critical component to predicting the performance of PULSAR. If the individuals sequenced within the pedigree are not predicted to share a high percentage of alleles IBD one should consider alternative phasing approaches, such as those designed for singletons.

Our simulations, using realistic genotypes and chromosomal haplotypes, show that PULSAR is fairly robust to genotyping errors. In fact PULSAR can be used to correct some genotyping errors. PULSAR “fixes” incorrect genotypes using a rudimentary method, a simple majorities vote among individuals sharing a haplotype. A weighting scheme based on the confidence of genotype calls or based on the number of reads supporting a given allele call may improve PULSAR’s ability to fix genotype errors.

We have not investigated the impact of errors in the pedigree structures themselves on PULSAR (including the existence of unknown relationships between pedigree founders). We advocate that pedigree relationships are confirmed or estimated analytically(Sun et al. 2002; Sun and Dimitromanolakis 2014) before one embarks on efforts to establish phase, correct genotyping errors, or impute missing genotypes.

PULSAR was designed specifically to work on whole genome sequence data, based on the rationale that the vast number of rare sequence variants in the human genome would produce a high density of LSAs in pedigrees. While whole genome sequencing clearly is the preferred way to characterize the genome, at the present time many studies only involve exome sequencing data or dense SNP genotyping, mainly due to cost. We have not investigated the performance of PULSAR on these other genotyping platforms. SNP genotyping panels are biased towards common SNPs. For that reason, we anticipate that PULSAR would be of limited utility for such data in extended pedigrees, where only very few common SNPs would be introduced only once into a given pedigree and thus serve as LSAs to the algorithm. On small pedigrees with relatively few founders we expect that the PULSAR algorithm would be much less impacted and work quite well.

There are a number of potential improvements and extensions to PULSAR that we could present, but these are beyond the scope of this manuscript. However, we did explore the utility of pre-screening variants upfront based on allele frequency when identifying putative LSAs. When pedigree founders are not available for sequencing, which will often be the case in reality, then such filtering based on minor allele frequency was shown to be a useful procedure for reducing the number of false positives among putative LSAs and thus improve the performance of the method. However, allele frequency estimates should be reliable to be effective as pre-filters. They can be taken either from the study at hand, if there are sufficiently many founders, or taken from reference panels. If one takes the estimates from reference panels, one should take care that the ethnicities be identical (or as close to it as possible), and that the sequencing quality be high and, ideally, based on the same technology platform and methods.

The algorithm that we have presented is a rules-based procedure, rather than a maximum likelihood approach. Conceptually, there are many methods that can be used to infer phase in pedigrees using maximum likelihood. For example, one could use the Elston-Stewart algorithm(Elston and Stewart 1971), as implemented in LINKAGE(Lathrop et al. 1984) or FASTLINK(Cottingham et al. 1993), to infer phase. The practical difficulty is that the Elston-Stewart algorithm is very limited in the number of variants that can be jointly analyzed, which would make it necessary to break the genome up into a vast number of overlapping segments, whose phased haplotype data would then have to be patched together. Alternatively, the Lander-Green algorithm(Lander and Green 1987), as implemented in the software package MERLIN(Abecasis et al. 2002), can analyze many variants simultaneously, but are limited in the number of individuals that can be handled. For large pedigrees, this would then necessitate breaking them into overlapping subcomponents, followed by re-combining the phased genotypes. These are not trivial tasks, since joining marker segments of pedigree fragments may lead to Mendelian inconsistencies and other problems. To overcome some of the limitations of the Elston-Stewart algorithm and the Lander-Green algorithm, a number of Monte Carlo Markov Chain methods have been developed, including LOKI(Heath 1997) and SimWalk2(Sobel and Lange 1996). However, these methods are extremely computationally intensive and are not readily applicable on whole genome sequence data. All of these methods are also impacted by genotyping errors and missing genotype data to varying degrees, and are no panacea.

There are also a number of methods for phasing of singleton individuals(Browning and Browning 2011), which are often based on hidden Markov models, such as BEAGLE(Browning and Browning 2007). Currently, the most prominent phasing method for singleton individuals are variants of the approximate coalescent models, such as SHAPEIT(Delaneau et al. 2011) and MACH(Li et al. 2010). Their accuracy and computational speed has greatly improved recently, aided by the availability of ever-larger reference panels for some of the major ethnic groups. The problems with applying these methods to pedigrees is that suitable reference panels are not presently available in many smaller populations, which are particularly suitable for pedigree studies. Without large reference panels, statistical phasing methods perform much worse. In addition, treating family members as unrelated individuals during phasing will often result in phased data that is inconsistent with Mendelian rules of inheritance. As part of our research for this paper, we have investigated the utility of duoHMM(O’Connell et al. 2014), which is implemented within the program SHAPEIT2(Delaneau et al. 2011), which seeks to correct haplotypes produced by statistical phasing using the restrictions on inheritance observed in duos. However, when applied to the simulations described in this study this method frequently failed to run on most of the simulated pedigree structures, and thus we chose to exclude it from the benchmarking results that were presented.

Given the different weaknesses and limitations of the various alternative phasing methods when applied to pedigree data, there appears to be a need for a rational method of combining phase results from multiple methods, especially those with different premises. Results from a combination of methods may prove to be more accurate and complete, yet also practical. For example, the phasing output from PULSAR could be used as input for Monte Carlo Markov Chain methods, providing an initial plausible phased genotyping state that could then be refined by MCMC. Or the output from statistical phasing for singletons may be taken as input for pedigree-based phasing methods. In this way, one could harness the high accuracy of methods that directly observe inheritance within pedigrees with statistical phasing methods that are capable of inferring phase in segments of the genome where inheritance is not directly observable given the individuals that are sequenced.

Here we have presented a novel algorithm, PULSAR, with associated software, for phasing WGS genotypes in a broad range of pedigree sizes and structures. The high accuracy of the haplotypes produced by the PULSAR algorithm, which often accurately span entire chromosomes, is promising. The approach may also be a useful tool for genotype error checking and correction. Based on evidence from our benchmarking we conclude that PULSAR is a suitable and practical phasing approach across a broad range of pedigree structures when a reasonable proportion of the pedigree members are sequenced.

## Methods

### Pedigrees used for simulation studies

We chose to investigate the phasing and imputation accuracy of PULSAR via simulation in a variety of pedigree sizes and structures. Nuclear pedigree structures were composed of two parents and a range of 1 to 7 children. As a variance reduction technique in our simulations, each smaller pedigree was generated as a subset of the larger pedigrees. In other words, the pedigree with 1 child is a subset of the pedigree with 2 children, and so on. Seven larger, multigenerational pedigree structures were chosen from the San Antonio Mexican American Family Studies(Mitchell et al. 1996; Hunt et al. 2005) (SAMAFS), having 11,14,20,28,55,78 and 94 individuals comprising 3,4,4,4,6,6 and 5 generations, respectively. In these pedigrees, 4,3,3,16,29,31, and 61 individuals are sequenced, respectively. Diagrams of these pedigrees, generated using Cranefoot(Makinen et al. 2005), are included in the supplementary materials. Altogether, 14 different pedigree structures were chosen to represent a broad range of sizes and structures in order to investigate the accuracy and completeness of the phasing results. Two additional pedigree structures were simulated for the purpose of investigating computational scalability of the software: pedigrees with two parents and a range between 1 and 1000 children, and pedigrees in which each generation consisted of one offspring of the previous generation and one married-in founder for 2-50 generations.

### Simulation of whole genome sequencing data

Whole genome sequencing data for 84 male X chromosomes (excluding the pseudoautosomal regions) from British (GBR) and Finish (FIN) populations from the 1000 Genomes Project(Genomes Project et al. 2015) were used as the set of potential founder chromosomes in the simulation studies. Utilizing real male X chromosomes has the advantage that chromosomal haplotypes are known while also maintaining the complexities of real whole genome sequencing data, such as the minor allele frequency distribution of variants, linkage disequilibrium, and the existence of small IBD segments shared between distantly related individuals(Browning and Browning 2007), though the X chromosome may differ slightly in these characteristics from the autosomes. After filtering out pseudoautosomal regions and loci with more than 2 alleles, there were a total of 318,912 polymorphic variants in the seed dataset. These seed chromosomes were shortened in simulation to 200,000 variants in order to accommodate limitations in the software package AlphaPhase1.1(Hickey et al. 2011), by considering a smaller section of the chromosome. Alphaphase1.1 was used for benchmarking because it implements the “long range phasing” approach in pedigrees (see below), although it was not designed for phasing WGS data. Within each pedigree, founder chromosomes were sampled from these 84 chromosomes randomly without replacement, which assumes that the pedigree founders represent a random population sample. Inheritance of haplotypes within the pedigrees was simulated by “gene dropping”, utilizing a probability of recombination between variants that corresponds to 1 recombination every 100 Mb, a rough estimate of the actual recombination rate in humans. Chromosomes not used as pedigree founder chromosomes were used as a reference panel from which to calculate minor allele frequency. In summary, for individuals in these pedigrees we simulated genotypes and chromosomal haplotypes with many of the characteristics of real data.

### Whole genome sequencing data

Whole genome sequencing has been generated for many participants in the SAMAFS. SNV and di-allelic INDELs with at least 5 observations of the minor allele were homogenized and merged from vendor provided (Complete Genomics, Illumina) sequencing genotype calls from 2330 directly sequenced genomes, resulting in 27,160,796 genetic variants. This data is available through dbGaP Study Accession: phs000462.v1.p1. Chromosome 21 (356,545 variants) was selected for analysis. Individuals and variants with genotyping rates below 99% were excluded, resulting in 354,466 variants. We chose to focus on the same 7 pedigrees of the SAMAFS for both real and simulated datasets in order to maximize the comparability of the real-world data to the simulated data. The 7 pedigrees are comprised of three hundred individuals, of which 147 are sequenced.

### Benchmarking

We set out to benchmark PULSAR against long range phasing (LRP) (Kong et al. 2008), arguably the current best approach for phasing genotypes in multigenerational pedigrees(Browning and Browning 2011). AlphaPhase1.1(Hickey et al. 2011) is an implementation of the LRP algorithm pioneered by Kong *et al*(Kong et al. 2008), which phases haplotypes using the assumption that the genotype data of individuals sharing a haplotype IBD must share at least one allele identical-by-state (IBS) at each locus in the shared region. Functionally, the method searches for opposing homozygous alleles, which excludes two individuals from sharing a segment IBD at that genomic location (assuming no mutation or genotyping error). LRP is highly accurate when at least one individual sharing the IBD segment is homozygous for a given variant.

We utilized three metrics for the performance comparison in the simulations: switch error rate(Lin et al. 2002) (SER) to assess phasing accuracy, the proportion of heterozygous genotypes that are phased, and the ‘time’ function in the Linux environment to assess the run time of the software. SER measures the rate in which adjacent heterozygous genotypes are phased incorrectly with respect to each other.

## Data Access

PULSAR is available at <u>https://github.com/AugustBlackburn/PULSAR_1.0</u>.

## Acknowledgements

We would like to thank the participants of the SAMAFS. The SAMAFS whole genome sequence data were obtained as part of the T2D-GENES Consortium, which is supported by NIH grants U01 DK085524, U01 DK085584, U01 DK085501, U01 DK085526, and U01 DK085545. The San Antonio Family Heart Study and San Antonio Family Diabetes/Gallbladder Study provided pedigree collection data, which are supported by NIH grants R01 HL0113323, P01 HL045222, R01 DK047482, and R01 DK053889. This work was partially funded by NIH grant DK099051.

